# OPTIMIR, a novel algorithm for integrating available genome-wide genotype data into miRNA sequence alignment analysis

**DOI:** 10.1101/479097

**Authors:** Florian Thibord, Claire Perret, Maguelonne Roux, Pierre Suchon, Marine Germain, Jean-François Deleuze, Pierre-Emmanuel Morange, David-Alexandre Trégouët, on behalf of the GENMED Consortium

## Abstract

Next-generation sequencing is an increasingly popular and efficient approach to characterize the full set of microRNAs (miRNAs) present in human biosamples. MiRNAs’ detection and quantification still remain a challenge as they can undergo different post transcriptional modifications and might harbor genetic variations (polymiRs) that may impact on the alignment step. We present a novel algorithm, OPTIMIR, that incorporates biological knowledge on miRNA editing and genome-wide genotype data available in the processed samples to improve alignment accuracy.

OPTIMIR was applied to 391 human plasma samples that had been typed with genome-wide genotyping arrays. OPTIMIR was able to detect genotyping errors, suggested the existence of novel miRNAs and highlighted the allelic imbalance expression of polymiRs in heterozygous carriers.

OPTIMIR is written in python, and freely available on the GENMED website (http://www.genmed.fr/index.php/fr/) and on Github (github.com/FlorianThibord/OptimiR).

## Introduction

With an average length of 22 nucleotides, microRNAs (miRNAs) belong to a class of small non-coding RNAs known to regulate gene expression by binding messenger RNAs (mRNAs) and interfering with the translational machinery (Bartel, 2004; Filipowicz et al., 2008). MiRNAs are transcribed from primary miRNA sequences (pri-miRNAs) and fold into a hairpin-like structure, which is sequentially processed by two ribonucleases, DROSHA and DICER. The former cleaves the pri-miRNA into a pre-miRNA and the latter completes the miRNA’s maturation by cleaving the pre-miRNA near its loop to produce a miRNA duplex composed of 2 mature strands (Kim et al., 2009). Exceptionally, some miRNAs follow a slightly different pathway where only one ribonuclease is needed to complete the maturation (Kim et al., 2016). In any case, only one of the two mature strands is loaded in an effective protein complex called RISC, while the other is degraded (Kawamata and Tomari, 2010). This selection seems mostly driven by the thermodynamic stability of both ends forming the duplex (Meijer et al., 2014).

There is emerging interest in performing miRNA profiling in body fluids or tissues in order to identify novel molecular determinants of human diseases (Mitchell et al., 2008; Pulcrano-Nicolas et al., 2018). Such miRNA profiling can be achieved using either hybridization (microarray), Next Generation Sequencing (NGS) or Real Time-quantitative Polymerase Chain Reaction (RT-qPCR) techniques. With 2,588 known mature miRNAs in humans according to miRBase version 21 (Kozomara and Griffiths-Jones, 2014), RT-qPCR would be cumbersome on a genomic scale, but is widely recognized as a gold standard for the validation of few miRNAs. The NGS technology is becoming more popular than microarrays because of its greater detection sensitivity, and higher accuracy in differential expression analysis (Git et al., 2010; Tam et al., 2014). NGS applied to small RNAs revealed a great diversity in the sequences of mature miRNAs originating from the same hairpin. This diversity is mostly attributable to the deletion and addition of nucleotides at the miRNAs’ extremities (also known as trimming and tailing events, respectively), due to the activity of terminal nucleotidyl transferases, exoribonucleases, or imprecise cleavage by DROSHA and DICER (Wyman et al., 2011; Neilsen et al., 2012; Ameres and Zamore, 2013). To a lesser extent, the ADAR protein acting on double stranded RNAs and responsible for A-to-I editing is also known to target miRNAs (Nishikura, 2016). These post-transcriptional editing mechanisms have been shown to affect miRNAs’ function and stability (Kawahara et al., 2007; Chiang et al., 2010; Burroughs et al., 2010; Katoh et al., 2015). Lastly, genetic variations have also been shown to contribute to the sequence diversity of miRNAs, and to affect their function and expression (Mencia et al., 2009; Gong et al., 2012; Han and Zheng, 2013; Cammaerts et al., 2015). MiRNAs subject to post-transcriptional events and/or genetic variations are generally referred to as isomiRs. In the following, we will use the expression “polymiR” to refer to the subclass of isomiRs harbouring genetic polymorphisms in their miRNA sequence.

The first step in the bioinformatics analysis of miRNA sequencing (miRSeq) data consists in aligning sequenced reads to a reference library of mature miRNAs. This step may be challenging because 1 - the aforementioned variability of isomiRs could lead to imperfect alignments to the reference library; 2 - sequenced reads may correspond to (fragments of) other molecules (e.g. other small non coding RNAs like piRNA, tRNA, yRNA, …), captured during the preparation of the libraries, that might share a high similarity with miRNAs because of their small length and thus might be confused with miRNAs (Chen and Heard, 2013; Heintz-Buschart et al., 2018); 3 - some miRNAs have homologous sequences that are identical or very similar, thus a single read might align ambiguously to multiple reference sequences. In this work, we investigate the impact of the presence of polymorphisms in the sequence of mature miRNAs on their alignment and their expression in the context of miRSeq profiling applied to samples that have also been typed for genome-wide genotype data. A situation we anticipate to become rather common with the rise of increasingly affordable genome-wide association studies (GWAS) and the decreasing cost of next generation exome/genome sequencing techniques. In that context, we developed an original bioinformatics workflow called OPTIMIR, for pOlymorPhism inTegratIon for MIRna data alignment, that integrates genetic information from genotyping arrays or DNA sequencing into the miRSeq data alignment process with the aim of improving the accuracy of polymiRs alignment, while accommodating for other isomiRs detection and ambiguously aligned reads. In addition, OPTIMIR allows to assess the association of genotypes on polymiRs with corresponding polymiRs’ expression. OPTIMIR was evaluated in the plasma samples of 391 individuals, part of the MARTHA study (Oudot-Mellakh et al., 2012).

## Results

OPTIMIR is composed of three main steps (Materials and Methods). First, miRSeq data are aligned to a reference library upgraded with sequences integrating alternative alleles of genetic variations. A correction is then applied for ambiguous and unreliable alignments via a scoring approach. Finally, polymiR alignments are evaluated to only retain those that are consistent with input genotypes in case these have been provided by the user. The general workflow was summarized in Figure 1.

**Figure 1:**
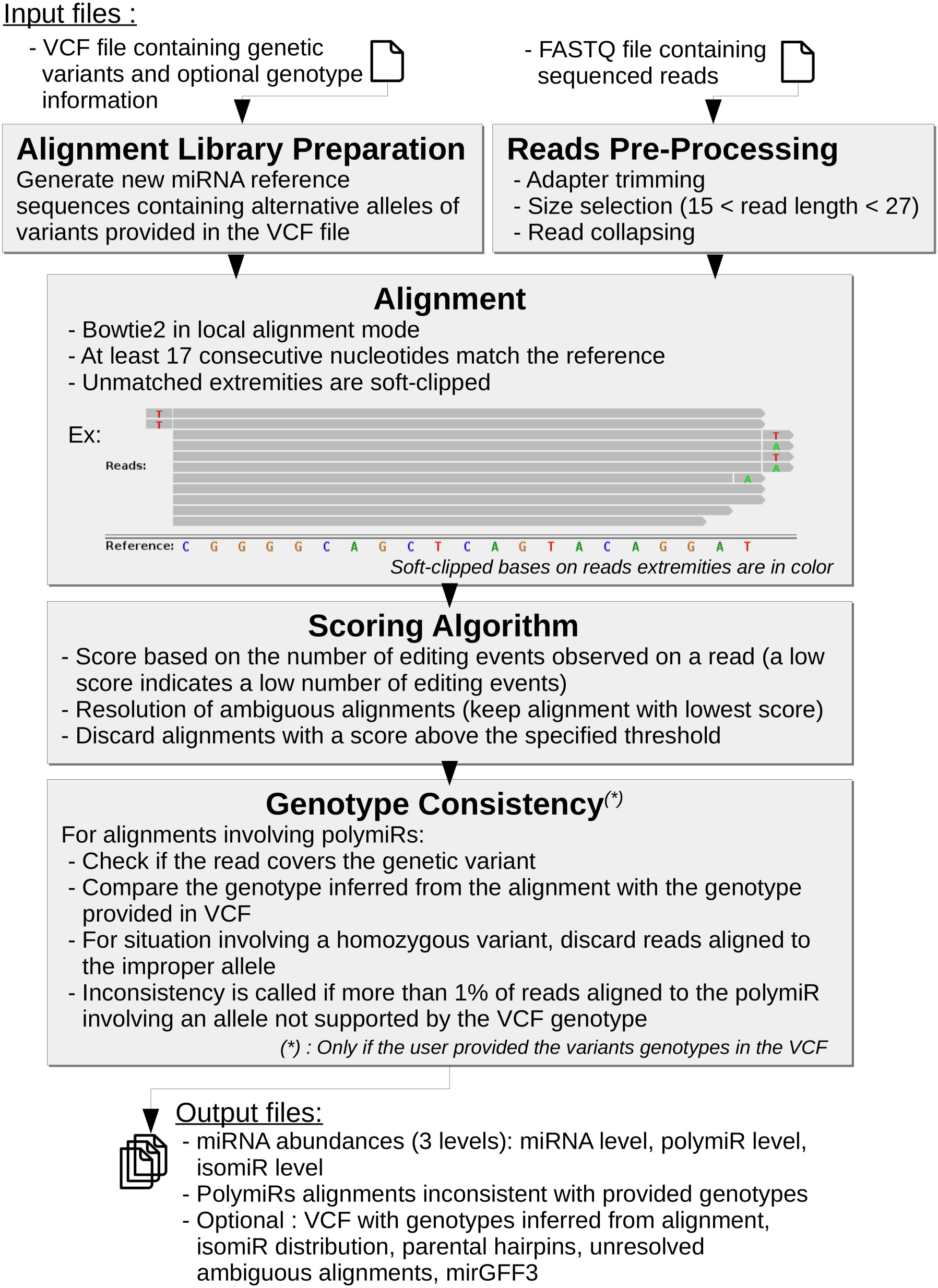
OPTIMIR workflow. From an optional VCF file and a FASTQ file, OPTIMIR performs the alignment of miRSeq data and results in the generation of abundances files containing the expressions of sequenced miRNAs.

### MiRNA alignment

OPTIMIR was evaluated on 391 miRNA sequencing data files totalizing 7,390,947,662 sequencing reads. After pre-processing the sequenced reads, that included adapters’ removal and selecting only reads with size ranging between 15 and 27 nucleotides, 2,922,446,965 reads (39.54% of total reads) remained for alignment. 562,040,494 of these reads (19.23%) were then mapped to mature miRNA reference sequences, of which 10,937,479 (1.95% of mapped reads) aligned ambiguously to two sequences or more. The application of the OPTIMIR scoring algorithm for alignment disambiguation resulted in a unique solution for 91.6% of these cross-mapping reads. OPTIMIR computes a score based on the number of editing events that a sequenced read could be compatible with and keeps alignments with the lowest scores (i.e the lowest number of editing events). For example, if a given read aligns to 2 reference sequences, perfectly on the first one (score 0), but with a missing base on the 3’ end (score 1) on the second, then only the first alignment is kept. A score of 0 corresponds to a perfect match, and each editing event adds penalties to the score. Modifications in the 5’ end are more penalizing, as they are less frequently observed. If a cross-mapping read receives an equal score on different alignments, then its weight is divided accordingly to the number of equivalent alignments (see Material and Methods). For 89.5% of reads with multiple alignments, the difference between the two lowest scores was greater than 2 (Figure 2.A) which would correspond to alignments that differ from each other by at least two modifications in the 3’ end.

**Figure 2:**
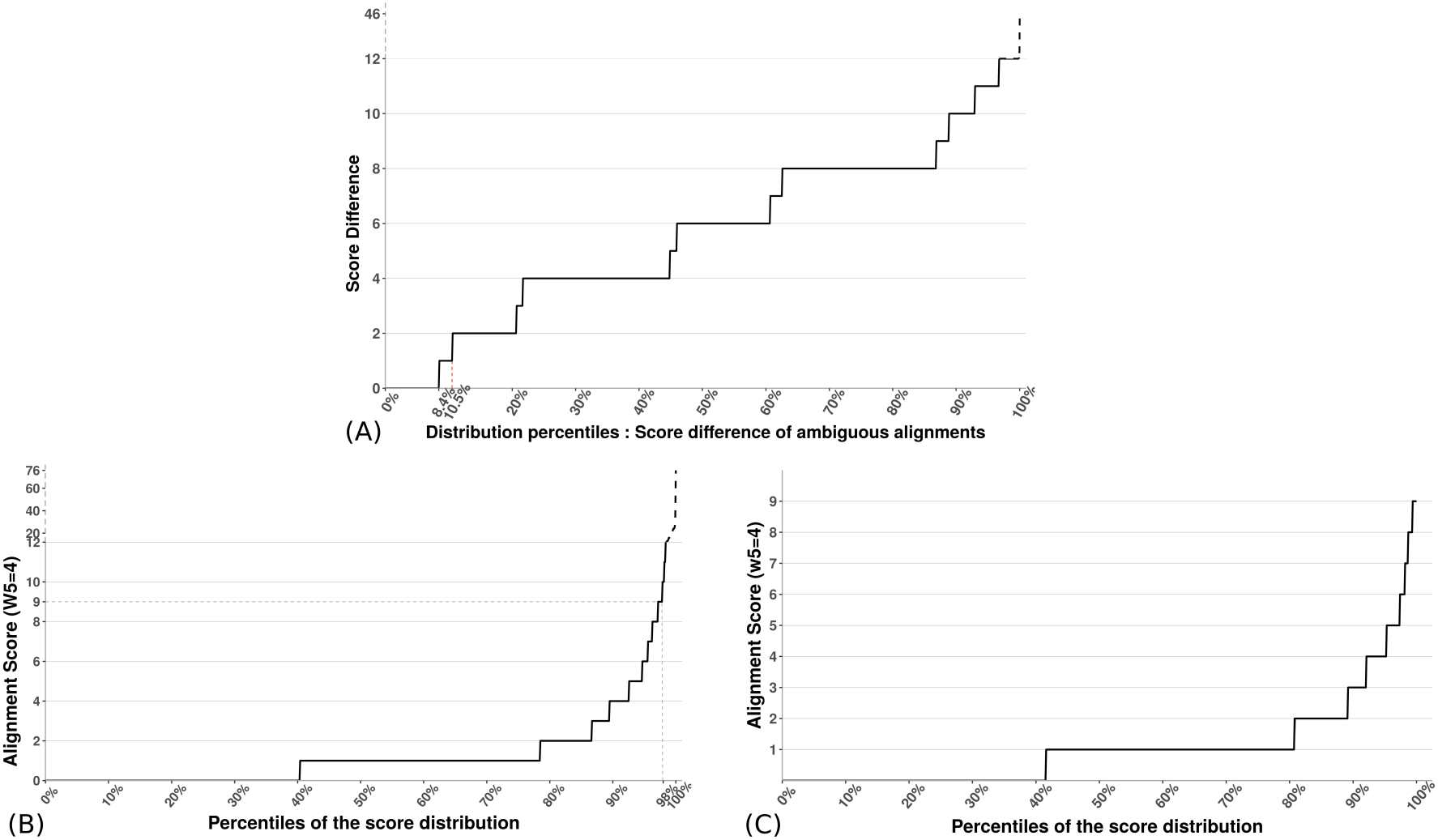
Influence of the scoring algorithm. (A) This figure represents the percentile distribution of the score differences between the two best alignments of each cross-mapping read. 8.4% of cross-mapping reads have a score difference of 0, which means they aligned on different sequences with the same score. These alignments could not be disambiguated. W5 indicates the penalty weight for events in the 5’ end, fixed to 4 in this study. (B) The alignment score percentile distribution after cross-mapping reads disambiguation represented in percentiles. 98% of alignments have a score lower or equal to 9. The remaining 2 % of alignments have a score ranging from 10 to 76, which were categorized as unreliable isomiRs because of the unlikely number of editing events they would have underwent. (C) The same distribution as in Figure 2.B but after removing alignments with a score higher than 9.

**Figure 3:**
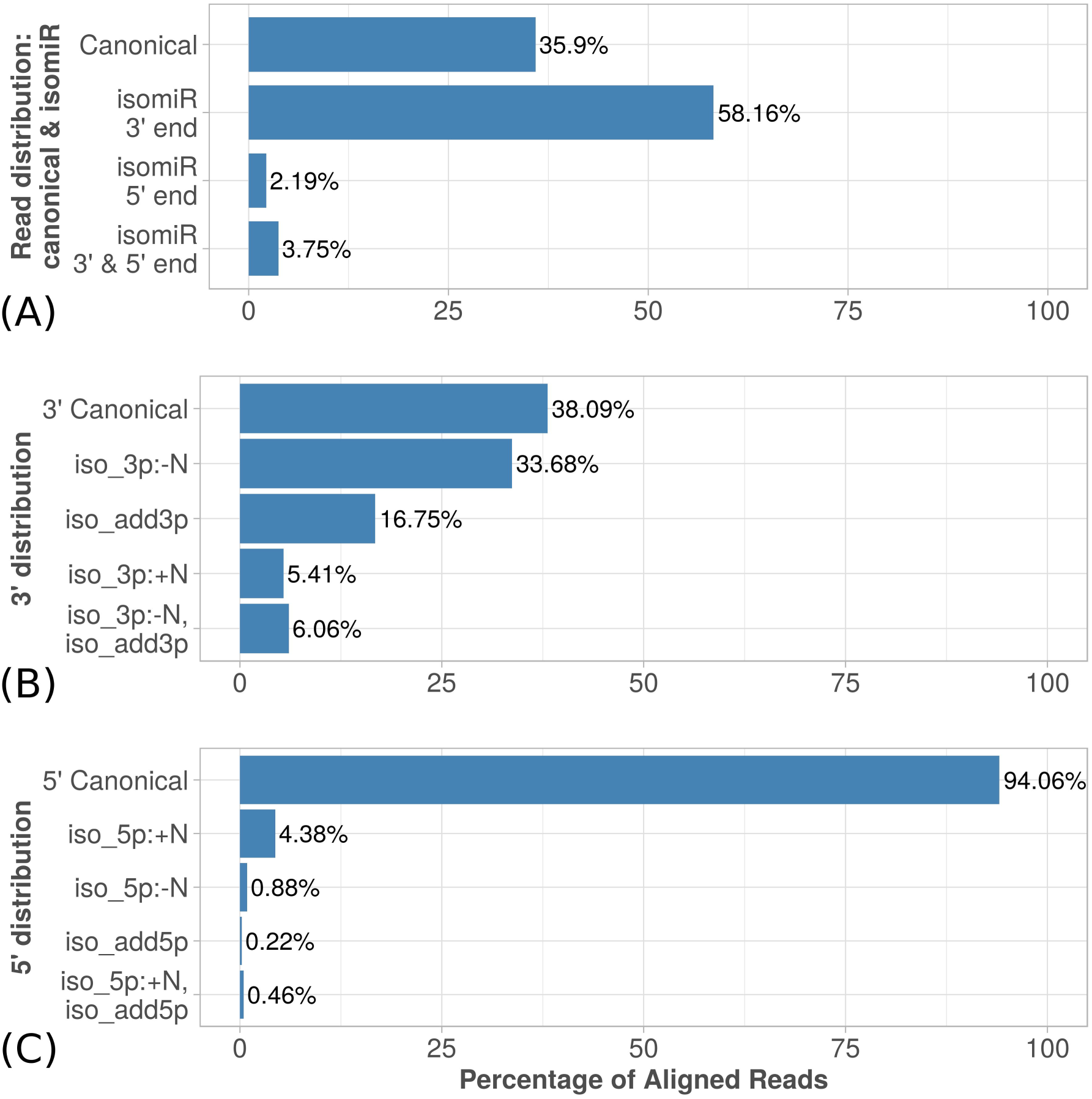
IsomiRs profiling. (A) Reads aligned by OptimiR distributed between: canonical miRNA (identical to the reference), isomiR 3’ end / 5’ end (modifications due to trimming or tailing observed on the 3’ or 5’ end, respectively), and isomiR with both ends edited. (B) Distribution of variations observed on the 3’ end of aligned miRNAs: 3’Canonical (no modification observed); iso_3p:-N (trimming); iso_3p:+N (templated tailing); iso_add3p (non templated tailing); iso_3p:-N,iso_add3p (combination of trimming and non templated tailing). (C) Distribution of variations observed on the 5’ end of aligned miRNAs: 5’Canonical (no modification observed); iso_5p:+N (trimming); iso_5p:-N (templated tailing); iso_add5p (non templated tailing); iso_5p:+N,iso_add5p (combination of trimming and non templated tailing).

After alignment disambiguation, scores ranged from 0 to 76 with 98.0% of alignment scores lower or equal to 9. Beyond this threshold, the quantile distribution curve rapidly increased indicating that scores with higher values are very sparse and suggesting that such alignments with a very high number of editing events are likely improper alignments (Figure 2.B). As a consequence, for the following, we decided to discard any alignment with a score greater than 9. Among the 550,946,055 remaining reads, more than 40% were perfectly aligned or involved templated additions, which were not penalized as these nucleotides are present in the parental hairpin sequence. An additional 40% of alignments involved reads with a single event in the 3’ end (see Figure 2.C).

### IsomiRs distribution

197,808,779 (35.9%) reads perfectly aligned to mature miRNAs, such reads being generally referred to as canonical miRNAs, and received an alignment score of 0. This confirms previous observations suggesting that a substantial amount of miRNAs are mainly represented by alternative isomiRs (Wu et al., 2018; Wallaert et al., 2017).

The most common observed editing events were on miRNAs’ 3’ end, with ~34% of trimming, ~17% tailing with non-templated nucleotides, ~5% tailing with templated nucleotides, and a similar proportion of trimming events followed by tailing. The latter modification could also be interpreted as nucleotide variation due to genetic variants, or other less frequent post transcriptional editing events such as A-to-I editing. It should also be noted that library preparation and sequencing could also contribute for a significant amount of fragment modifications that are falsely detected as isomiRs (Wright, 2019).

The 5’ end was much less frequently edited, with 94% of reads having no editing on this extremity. Nevertheless, the most frequently observed modification on 5’ ends was trimming that affected 4.4% of all reads. This may be of biological relevance since such trimming could shift the miRNA binding seed that is crucial for the miRNA to bind to its mRNA targets.

The distribution of 3’ and 5’ ends modifications on mapped reads observed over the 391 samples processed by OPTIMIR is shown in Figure 4.

**Figure 4:**
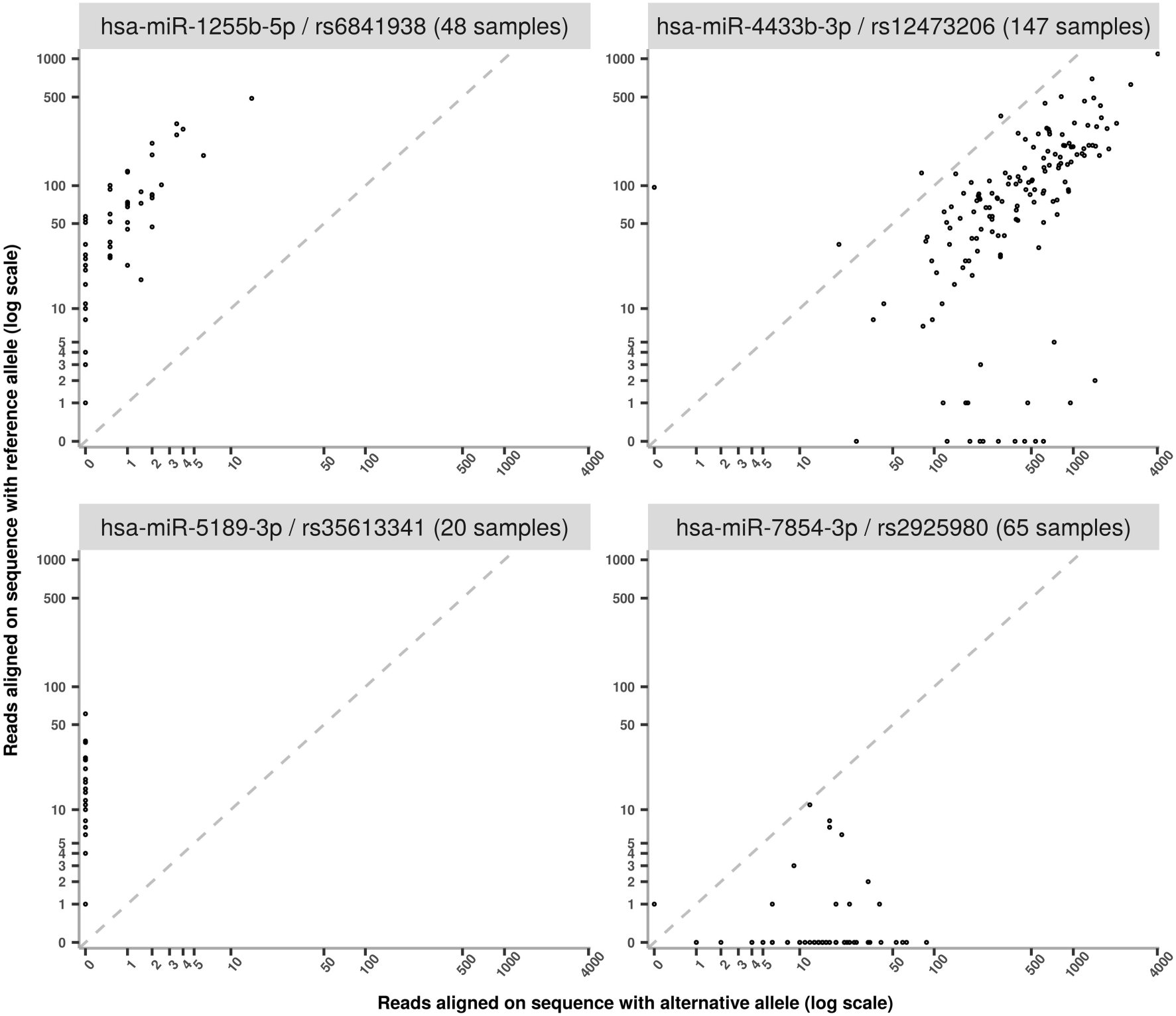
Examples of allele imbalanced expressions observed at 4 polymiRs. On the X-axis is shown the expression of the alternative allele while the Y-axis represents the expression of the reference allele. Expressions were log_10_-transformed.

### Alignment on polymiRs

Over all samples, 220,156 reads mapped to 46 polymiRs for a total of 1,786 distinct alignments. As detailed in Table 1, 19 polymiRs have reads that aligned to an alternative sequence that was introduced in the upgraded reference library. Two polymiRs (hsa-miR-6796-3p and hsa-miR-1269b) harbor two SNPs, and for both of them only the reference sequence with both common alleles were found to be expressed. Among the remaining 44 polymiRs that harbor only one SNP, 15 were expressed with both alleles. It is important to mention that the allele present in the miRBase reference sequence may not be the most common one (e.g. the rs2155248 and hsa-miR-1304-3p) which may lead to improperly discard reads if stringent alignment (with no mismatch allowed) to the original miRBase library is applied.

**Table 1:** Summary information at detected polymiRs.

| polymiR | rsID | Number of samples expressing the polymiR with genotype (1) |  |  | Number of reads aligned on polymiR sequence integrating the allele |  | Number of inconsistent reads removed on the sequence integrating the allele |  | Inconsistent alignments to investigate (2) |
| --- | --- | --- | --- | --- | --- | --- | --- | --- | --- |
|  |  | 0/0 | 0/1 | 1/1 | reference | alternative | reference | alternative |  |
| hsa-miR-4433b-3p | rs12473206 | 223 (/230) | 147 (/147) | 14 (/14) | 63 762 | 96 915 | 4 | 475 | 2 |
| hsa-miR-1255b-5p | rs6841938 | 244 (/323) | 48 (/66) | 2 (/2) | 29642 | 66 | 0 | 52 | 1 |
| hsa-miR-1304-3p | rs2155248 | 0 (/0) | 5 (/7) | 256 (/384) | 146 | 11 928 | 4 | 0 | 0 |
| hsa-miR-339-3p | rs72631820 | 213 (/386) | 3 (/5) | 0 (/0) | 6 293 | 0 | 0 | 21 | 1 |
| hsa-miR-7854-3p | rs2925980 | 22 (/159) | 65 (/167) | 35 (/65) | 392 | 2 015 | 1 | 4 | 0 |
| hsa-miR-4745-5p | rs10422347 | 86 (/300) | 14 (/81) | 0 (/10) | 2 330 | 15 | 0 | 0 | 0 |
| hsa-miR-5189-3p | rs35613341 | 42 (/196) | 20 (/161) | 0 (/34) | 1 315 | 0 | 2 | 1 | 0 |
| hsa-miR-7107-3p | rs3817551 | 20 (/157) | 33 (/177) | 0 (/57) | 1 104 | 1 | 122 | 0 | 5 |
| hsa-miR-4741 | rs7227168 | 40 (/293) | 4 (/93) | 0 (/5) | 695 | 0 | 0 | 0 | 0 |
| hsa-miR-548l | rs13447640 | 30 (/373) | 0 (/18) | 0 (/0) | 613 | 0 | 0 | 0 | 0 |
| hsa-miR-3620-5p | rs2070960 | 47 (/348) | 4 (/42) | 0 (/1) | 459 | 0 | 0 | 0 | 0 |
| hsa-miR-6826-5p | rs6771809 | 11 (/286) | 11 (/92) | 2 (/13) | 199 | 243 | 0 | 21 | 1 |
| hsa-miR-4638-5p | rs146528803 | 20 (/369) | 1 (/22) | 0 (/0) | 313 | 0 | 0 | 0 | 0 |
| hsa-miR-3622a-5p | rs66683138 | 6 (/231) | 4 (/138) | 0 (/22) | 241 | 24 | 0 | 0 | 0 |
| hsa-miR-4707-3p | rs2273626 | 4 (/86) | 6 (/195) | 0 (/110) | 162 | 0 | 0 | 0 | 0 |
| hsa-miR-1269a | rs73239138 | 5 (/236) | 2 (/131) | 0 (/24) | 107 | 31.5 (**) | 0 | 0 | 0 |
| hsa-miR-4781-3p | rs74085143 | 8 (/366) | 0 (/25) | 0 (/0) | 135 | 0 | 0 | 0 | 0 |
| hsa-miR-6763-3p | rs3751304 | 1 (/39) | 2 (/153) | 2 (/199) | 86 | 29 | 0 | 0 | 0 |
| hsa-miR-4804-5p | rs266435 | 0 (/7) | 3 (/91) | 7 (/293) | 0 | 114 | 1 | 0 | 0 |
| hsa-miR-4999-5p | rs72996752 | 5 (/189) | 1 (/163) | 0 (/39) | 86 | 26 | 0 | 1 | 0 |
| hsa-miR-6839-5p | rs7804972 | 1 (/153) | 2 (/196) | 0 (/42) | 56 | 0 | 1 | 0 | 0 |
| hsa-miR-4459 | rs73112689 | 3 (/245) | 1 (/133) | 0 (/13) | 51 | 0 | 0 | 0 | 0 |
| hsa-miR-585-3p | rs62376935 | 0 (/330) | 2 (/52) | 0 (/9) | 2 | 48 | 0 | 0 | 0 |
| hsa-miR-3928-5p | rs5997893 | 1 (/38) | 1 (/157) | 1 (/196) | 47 | 9 | 0 | 0 | 0 |
| hsa-miR-548al | rs515924 | 0 (/307) | 1 (/80) | 1 (/4) | 0 | 37 | 0 | 0 | 0 |
| hsa-miR-3117-3p | rs12402181 | 1 (/288) | 0 (/97) | 0 (/6) | 35 | 0 | 0 | 0 | 0 |
| hsa-miR-3130-3p | rs2241347 | 2 (/254) | 3 (/119) | 0 (/18) | 35 | 0 | 0 | 0 | 0 |
| hsa-miR-4532 | rs73177830 | 2 (/344) | 0 (/44) | 0 (/3) | 34 | 0 | 0 | 0 | 0 |
| hsa-miR-302c-3p | rs199971565 | 1 (/371) | 0 (/20) | 0 (/0) | 30 | 0 | 0 | 0 | 0 |
| hsa-miR-4772-5p | rs62154973 | 3 (/339) | 0 (/51) | 0 (/1) | 25 | 0 | 0 | 0 | 0 |
| hsa-miR-1343-5p | rs2986407 | 0 (/20) | 0 (/111) | 2 (/260) | 0 | 24 | 0 | 0 | 0 |
| hsa-miR-548ap-5p | rs4414449 | 2 (/47) | 11 (/180) | 1 (/164) | 16 | 4 | 4 | 0 | 0 |
| hsa-miR-6885-5p | rs78293125 | 1 (/341) | 0 (/47) | 0 (/3) | 15 | 0 | 0 | 0 | 0 |
| hsa-miR-548h-3p | rs73235381 | 3 (/379) | 0 (/11) | 0 (/1) | 14 | 0 | 0 | 0 | 0 |
| hsa-miR-4433a-5p | rs12473206 | 1 (/230) | 2 (/147) | 0 (/14) | 12 | 0 | 0 | 0 | 0 |
| hsa-miR-6887-5p | rs1688017 | 1 (/152) | 0 (/193) | 0 (/46) | 10 | 0 | 0 | 0 | 0 |
| hsa-miR-4482-5p | rs45596840 | 1 (/167) | 0 (/179) | 0 (/45) | 10 | 0 | 0 | 0 | 0 |
| hsa-miR-4520-5p | rs8078913 | 0 (/87) | 0 (/198) | 1 (/106) | 0 | 9 | 0 | 0 | 0 |
| hsa-miR-1229-5p | rs2291418 | 2 (/363) | 0 (/27) | 0 (/1) | 7 | 0 | 0 | 0 | 0 |
| hsa-miR-6805-3p | rs56312243 | 1 (/337) | 0 (/50) | 0 (/4) | 5 | 0 | 0 | 0 | 0 |
| hsa-miR-4302 | rs11048315 | 1 (/278) | 2 (/102) | 0 (/11) | 1 | 2 | 0 | 6 | 0 |
| hsa-miR-4520-3p | rs8078913 | 0 (/87) | 1 (/198) | 0 (/106) | 1 | 0 | 0 | 0 | 0 |
| hsa-miR-548h-5p | rs9913045 | 0 (/143) | 0 (/197) | 0 (/51) | 0 | 0 | 1 | 0 | 0 |
| hsa-miR-6863 | rs12708966 | 1 (/385) | 0 (/6) | 0 (/0) | 1 | 0 | 0 | 0 | 0 |
| hsa-miR-6796-3p | rs3745199 | 3 (/131) | 0 (/193) | 0 (/67) | 56 | 0 (*) | 0 | 0 | 0 |
|  | rs3745198 | 3 (/131) | 0 (/190) | 0 (/70) |  |  |  |  |  |
| hsa-miR-1269b | rs12451747 | 0 (/82) | 1 (/171) | 0 (/138) | 31.5 (**) | 0 (*) | 0 | 0 | 0 |
|  | rs7210937 | 1 (/326) | 0 (/60) | 0 (/5) |  |  |  |  |  |
| TOTAL |  | 1062 | 400 | 324 | 108574,5 | 111540,5 | 140 | 581 | 10 |

(1) Between parenthesis is the total number of samples carrying the variant with homozygous reference allele (0/0), heterozygous (0/1) or homozygous alternative allele (1/1)
(2) Alignments with a high number of inconsistent reads (higher than 1% of the sum of reads gathered by the polymiR sequences), which is incompatible with sporadic events replicating a genetic variant.
(\*) For multi variant polymiRs, reads did not map to any of the sequence integrating alternative haplotypes
(\*\*) The sequence of hsa-miR-1269a integrating the rs73239138-A allele is identical to the hsa-miR-1269b sequence. Hence each of the 63 reads that cross-mapped on both sequence received a count of ½ on each sequence.

In total, 155 alignments involving 721 reads (0.33% of reads mapped to polymiRs) were found inconsistent with the individual genotypes, among which 145 alignments were supported by less than 5 reads. These alignments were discarded and not further discussed as they most probably correspond to sporadic mechanisms that mimic genetic variations such as modifications induced during library preparation or sequencing (Wright, 2019), or uncommon post-transcriptional editing events (see Materials & Methods). The remaining 10 alignments, involving 507 reads spread over 10 samples and 5 polymiRs, are detailed in Table 2 and further investigated in the next section.

**Table 2:** Inconsistent genotypes unlikely due to sequencing errors

| polymiR | rsID | Alleles:<br>Reference,<br>Alternative | MAF | RSQ | SAMPLE | Genotype<br>from vcf | Counts<br>Reference | Counts<br>Alternative | SANGER<br>validation |
| --- | --- | --- | --- | --- | --- | --- | --- | --- | --- |
| hsa-miR-4433b-3p | rs12473206 | C/G | 0.23 | 0.99 | PVP28 | C/C | 84 | 333 | **G/C** |
| hsa-miR-4433b-3p | rs12473206 | C/G | 0.23 | 0.99 | SSP20 | C/C | 135 | 8 | N.A. |
| hsa-miR-339-3p | rs72631820 | T/C | 0.009 | 0.98 | CED24 | T/T | 139 | 7 | N.A. |
| hsa-miR-6826-5p | rs6771809 | T/C | 0.14 | 0.98 | DTD19 | T/T | 1 | 20 | N.A. |
| hsa-miR-1255b-5p | rs6841938 | G/A | 0.09 | 0.96 | AJA9 | G/G | 92 | 28 | **G/G** |
| hsa-miR-7107-3p | rs3817551 | T/G | 0.36 | 0.96 | MSM20 | G/G | 27 | 0 | N.A. |
| hsa-miR-7107-3p | rs3817551 | T/G | 0.36 | 0.96 | MCC20 | G/G | 15 | 0 | **G/G** |
| hsa-miR-7107-3p | rs3817551 | T/G | 0.36 | 0.96 | NFN22 | G/G | 23 | 0 | **G/G** |
| hsa-miR-7107-3p | rs3817551 | T/G | 0.36 | 0.96 | NHN19 | G/G | 25 | 0 | **G/G** |
| hsa-miR-7107-3p | rs3817551 | T/G | 0.36 | 0.96 | GLG20 | G/G | 17 | 0 | **G/G** |

MAF: Minor Allele Frequency
RSQ : imputation quality criteria (r2)

### Investigation of inconsistent genotypes

Situations where a substantial number of reads aligned inconsistently on a polymiR were reported by OptimiR to allow for further investigations.

The first case of inconsistent genotype concerns individual PVP28 imputed to be homozygous for the rs12473206-G allele while showing numerous reads mapping to both versions of the hsa-miR-4433b-3p polymiR. Sanger re-sequencing revealed that this individual is in fact heterozygous for this variant which is much more compatible with the number of observed reads at this locus. A rather similar inconsistent genotype was observed for individual SSP20 but lack of available DNA did not allow us to perform Sanger validation. Lack of DNA also prevented us from investigating deeper the inconsistent genotypes observed for rs72631820/hsa-miR-339-3p and rs6771809/hsa-miR-6826-5p.

The inconsistent genotype observed for individual AJA9 at rs6841938 and hsa-miR-1255-5p was challenging. Sanger re-sequencing confirmed that this individual was indeed homozygous for the rs6841938- G allele although it has numerous reads mapping to both versions of hsa-miR-1255-5p. However, the use of BLAST web-service from NCBI (Altschul et al., 1990) revealed that reads aligned on the polymiR sequence containing the rs6841938-A allele perfectly match to the chr1:167,998,699-167,998,720 region but on the opposite strand where hsa-miR-1255-3p should be located. This observation could be compatible with the presence on this opposite strand of an unreported miRNA locus as it has already been observed for other miRNAs (e.g: hsa-mir-4433a and hsa-mir-4433b). Going back to the discarded alignments revealed that 18 other samples homozygous for rs6841938-G had 1 or 2 reads that mapped to this opposite strand. These alignments had been discarded since they were supported by less than 5 reads (see *Genotype consistency analysis* section in the Material & Methods).

The last inconsistent genotypes were observed at rs3817551/hsa-miR-7107-3p for five independent individuals. For 4 of them with available DNA, Sanger re-sequencing confirmed the initial homozygous genotype for rs3817551-G allele while these individuals were found to express only the hsa-miR-7107-3p version with the T allele. On average, these inconsistent alignments received an alignment score of 1.41, and none of the reads involved mapped to other sequences. Concerning the samples that had reads aligned on this same reference with a consistent genotype, the average alignment score was very close to 1.47. This score indicates that reads share a high sequence similarity with the hsa-miR-7107-3p, and there is no significant difference between the group with an inconsistent genotype, and the group with a consistent one. BLAST analysis did not enable us to identify any homologous sequence that could explain these observations that still remain to be further investigated in order to be sure that associated reads do well originate from hsa-miR-7107-3p.

### Analysis of polymiRs allele specific expression

As shown in Table 1, 29 polymiRs with a SNP in their mature sequence were found to be expressed in individuals heterozygotes for this SNP. While one could anticipate that, in heterozygous individuals for such SNP, the polymiRs could have balanced expression (as measured by read counts) of both the reference and alternative sequences, this was hardly observed. Indeed, we observed a strong preference for either the reference or the alternative version of the polymiR in heterozygotes.

Figure 4 shows the alignments of 4 polymiRs involving heterozygous variants according to the vcf genotype. The dotted line represents a theoretical balanced expression between reference and alternative sequences. We can see that, for hsa-miR-1255b-5p/rs6841938 and hsa-miR-5189-3p/rs35613341, dots are close to the y-axis, which are situations where only the reference allele is expressed. The polymiR with the expression closest to allelic balance for many samples is hsa-miR-4433b-3p but, even then, the average rate of alternative reads across all 147 samples is of 0.8 (see Supplementary Table 4 for details on polymiRs involved in heterozygous situations). As the last case, 65 MARTHA individuals were heterozygous for the rs2925980 variant associated with polymiR hsa-miR-7854-3p. These genotypes could be considered as reliable as this SNP was directly typed on the array. Among these 65 individuals, 54 expressed only the alternative version of polymiR hsa-miR-7854-3p and the remaining 11 expressed both polymiR versions with a mean ratio of 0.96 in favor of the alternative allele.

Finally, we used the RNAfold program (Lorenz et al., 2011) to predict the secondary structure of the 29 pri-miRNA (i.e. hairpins) induced by the presence of a SNP in the polymiR sequence (Supplementary Figure S1). Most genetic variations create either a new bulge, or a wobble pairing, or have no impact on the secondary structure. A notable exception relates to rs35613341 located on the hsa-miR-5189-3p where the G allele completely changed the secondary structure of the hairpin (see Supplementary Figure S1-u) making it difficult to access for the DICER and DROSHA machinery. In MARTHA samples, 161 individuals were heterozygous and 34 homozygous for the rs35613341-G allele. None of these individuals were found to express the alternative sequence of polymiR hsa-miR-5189-3p which could support the hypothesis that the rs35613341-G allele impacts the maturation of this miRNA.

Lastly, by the completion of the OPTIMIR pipeline (with a scoring threshold of 9 as mentioned above and keeping only genotype consistent reads) on MARTHA samples, 7.45% of sequenced reads were aligned. This value had to be compared with 7.68% and 8.24% obtained by 2 other recent pipelines for miRSeq data, sRNAnalyzer (Wu et al., 2017) and miRge (Baras et al., 2015), respectively, executed using default parameters. These discrepancies could be explained by the different alignment strategies implemented by these tools, which by default allow up to 2 mismatches for read alignment, while OPTIMIR does not allow any mismatch in aligning sequenced read and rely on local alignment to capture isomiRs with modified extremities (see OPTIMIR description). When we ran sRNAnalyzer with only 1 mismatch allowed, the percentage of aligned reads dropped to 7.38% (compared to 7.68% with two mismatches). Of note, it was not possible to modify the number of authorized mismatches in miRge.

The files generated by OPTIMIR include: 1/ global abundances of miRNAs (counts of isomiRs and polymiRs are merged with the reference mature sequences’ counts); 2/ specific abundances for each polymiR sequence; 3/ specific abundances for each isomiR sequence; 4/ alignments that are inconsistent with provided genotypes; 5/ two annotation files containing details on templated nucleotides and alignments that could not be disambiguated. In addition, OPTIMIR allows to generate its results into the recently developed mirGFF3 format (Desvignes et al., 2018) aimed at unifying results of any miRSeq data analysis.

## Discussion

In this work, we propose a novel algorithm, OPTIMIR, for aligning miRNA sequences obtained from next generation sequencing and we applied it to plasma samples of 391 individuals from the MARTHA study. Borrowing some ideas from other alignment pipelines such as the addition of new sequences to the reference library corresponding to allelic versions of polymiRs (Baras et al., 2014; Russell et al., 2018) and a scoring strategy for handling cross-mapping reads (Urgese et al., 2016) or for discarding unlikely isomiRs (Bofill-De Ros et al., 2018), OPTIMIR has two features that make it unique. First, OPTIMIR is based on a scoring strategy that incorporate biological knowledge on miRNA editing to identify the most likely alignment in presence of cross-mapping reads. Second, OPTIMIR allows the user to provide genotype information, in particular data obtained from genome wide genotyping arrays, to improve alignment accuracy. This option revealed several interesting observations when OPTIMIR was applied to MARTHA plasma samples.

First, it allowed to identify improperly imputed genotypes despite overall good imputation quality. Second, it suggested the existence of a new miRNA not indexed in the miRBase v21 database that would be located on the opposite strand of the hsa-miR-1255b-5p. Thirdly, it suggested that reads aligned to hsa-miR- 7107-3p are likely false alignments and would more likely come from other non-coding fragments that share sequence similarity with hsa-miR-7107-3p. These last two hypotheses would need to be further validated but this is out of the scope of the bioinformatics workflow described in the current work. Even more interesting was the study of polymiRs’ expressions in heterozygous individuals for SNPs on these polymiRs. OPTIMIR clearly showed that plasma allelic expression of polymiRs is unbalanced for most polymiRs. One allelic version of a polymiR is much more expressed than the other and this is not necessarily the one carrying the most common allele. This observation is consistent with previous works showing that SNPs in (pri-) miRNA sequences can influence miRNA expression through their impact on the DROSHA/DICER RNAses’ machinery (Duan et al., 2007; Cammaerts et al., 2015). Epigenetic mechanisms could also explain the allelic imbalance expression of miRNAs (Morales et al., 2017).

Several limitations shall however be acknowledged. OPTIMIR requires to fix two parameters, a weight *W_5_* for penalizing 5’ end editing event (set to 4 in the current application) and a score threshold (set to 9 here) to discard unreliable alignments. The former tends to have little impact on the general findings (see Supplementary Table 2) while the second may be study-specific and may depend on the study sample-size and the kind of tissue analyzed. These parameters can be easily modified by the user. There is so far no gold standard program for miRNA alignment analysis but our preliminary study suggests that OPTIMIR aligns slightly less reads than two other recently proposed software, sRNAnalyzer (Wu et al., 2017) and miRge (Baras et al., 2015). This is likely due to the higher number of mismatches allowed by the laters for aligning reads while OPTIMIR tends to be more stringent. Without extensive investigations including experimental validations, it is not possible to really appreciate which alignments are the correct ones.

Finally, several improvements could be considered such as: 1/ the integration of A-to-I editing events in the definition of our reference library and of our scoring strategy, even if we anticipate that it might be difficult to distinguish these rare events from sequencing errors and 2/ the extension of the OPTIMIR workflow to analyze other small coding RNAs (e.g. piRNA and tRNA, rRNA, snRNA or yRNA derived fragments) that are generally sequenced together with miRNAs in a miRseq profiling. Applications to other tissues from the samples processed for miRSeq data deserve to be conducted to generalize the findings observed in the current plasma samples.

## Materials & Methods

### The MARTHA dataset

The MARseille Thrombosis Assocation study is a collection of patients with venous thrombosis (VT) recruited at the La Timone Hospital (Marseille, France) between 1994 and 2005 and aimed at identifying novel molecular determinants for VT and its associated endophenotypes (Oudot-Mellakh et al., 2012; Dick et al., 2014).

For the present study, 391 MARTHA participants with available plasma samples were processed for plasma miRNA profiling through miRSeq. These individuals had been previously typed for genome-wide genotyping arrays and imputed for single nucleotide polymorphisms (SNPs) available in the 1000G reference database (Germain et al., 2015).

MiRNA extraction and preparation followed the same protocol as the one previously described in (Roux et al., 2018). Briefly, from 400µL of plasma, total RNA was first extracted using the miRNeasy serum/plasma kit for Qiagen. MiRNA libraries were then prepared using the NEBNext Multiplex Small RNA Library Preparation Set for adapter ligation and PCR, with adapter sequences GATCGGAAGAGCACACGTCTGAACTCCAGTCAC (3’ adapter) and CGACAGGTTCAGAGTTCTACAGTCCGACGATC (5’ adapter) followed by a size selection using AMPure XP beads. Pools of equal quantity of 24 purified libraries were constructed and tagged with different indexes. Pools were then sequenced using a 75bp single-end strategy on an Illumina NextSeq500 instrument.

### The OPTIMIR workflow

#### Alignment

##### Pre-alignment data processing

3’ adapters were removed using cutadapt (Martin, 2011) with a base quality filter set to 28. Remaining reads with sequence length between 15 and 27, which generally correspond to miRNA sequences, were then kept for alignment. Note that identical reads were collapsed together to decrease computational burden associated with processing *n* times *n* identical reads.

##### Definition of an alignment reference library

Read alignment generally starts by the selection of a reference library to which reads shall be aligned. The miRBase 21 database (Kozomara and Griffiths-Jones, 2014) containing known human mature miRNAs is usually adopted for miRSeq data. We first upgrade this reference library by adding new sequences corresponding to the alternate forms of polymiRs showing genetic polymorphisms in their mature sequence as previously proposed (Baras et al., 2014; Russell et al., 2018). In case a polymiR contains more than one polymorphism, new sequences corresponding to all possible haplotypes are generated. These variants, that are provided by the OPTIMIR’s user in a vcf format file (Danecek et al., 2011), are mapped to miRNAs using miRBase miRNA coordinates file (i.e. positions of miRNAs on the human reference genome). The generation of new sequences is automated via a standalone python script provided with the OPTIMIR pipeline.

For the current application to MARTHA GWAS data, we identified 88 single nucleotide polymorphisms (SNPs) for which we have a reliable genotype data, defined as SNPs with imputation r^2^> 0.8. Note that some SNPs may map to 2 distinct miRNAs if the latter are transcribed from opposite strands. Some miRNAs may also have more than one SNP in their sequence. In our application, 5 SNPs mapped to 2 distinct miRNAs and 3 miRNAs contained more than one SNP (see Supplementary Figure 2 for examples). As a result, the reference library was upgraded with 96 new alternative sequences corresponding to all possible haplotypes derived from the 90 identified polymiRs.

##### Read alignment process

For read alignment, we opted for the bowtie2 software (Langmead and Salzberg, 2012) that can handle trimming and tailing events at the reads’ extremities via its local alignment mode which has been shown to be efficient for miRSeq data alignment (Ziemann et al., 2016). Only reads with a sequence of at least 17 consecutive nucleotides (defined as the alignment seed) that perfectly match with the reference library are kept in the analysis (see Supplementary Table 1 for details concerning the choice of the seed value and its consequences on isomiR detection). In summary, the OPTIMIR pipeline does not authorize any mismatch in the central sequence of a read but allows variations at its extremities to address post-transcriptional editing. The handling of miRNAs with genetic variations is addressed by the use of the upgraded reference library described above. For miRNAs that underwent tailing events, or trimming events followed by tailing events, additional bases exceeding or differing from the reference are soft-clipped and do not participate in the alignment. With a limited read length of 27 nucleotides, the maximum number of bases that can be soft-clipped was set to 10. Reads were allowed to align to multiple reference sequences in order to take into account the different mature miRNAs with similar sequences from which they could originate. Finally, we did not allow reverse complement alignment as small RNAs were first ligated with different 5’ and 3’ adapters before single-end sequencing, which implies that RNA strands were sequenced in only one direction.

#### Resolution of ambigous alignments

To handle multiple ambiguous alignments, OPTIMIR integrated a scoring algorithm aimed at identifying the most plausible alignment(s) while discarding likely erroneous ones. Of note, beforehand, for reads mapping to a mature miRNA that can be produced by two different pri-miRNAs (e.g. hsa-miR-1255b-5p can originate from hsa-mir-1255b-1 or hsa-mir-1255b-2 located on chromosome 4 and 1, respectively), we used the information on templated tailed nucleotides (i.e. nucleotides in the pri-miRNA sequences (also available in the miRBase 21) that surround the mature miRNA sequence) to deduce from which locus these reads might come from. This information, that has no impact on the alignment per se, is stored in an output file (named *expressed_hairpins.annot*).

Each alignment was assigned a score based on the number of trimming and tailing events that could make a given read perfectly match with a mature miRNA sequence. Since trimming and tailing are frequently observed in the 3’ end of miRNAs but rare in the 5’ end (Neilsen et al., 2012; Wu et al., 2018), a more penalizing weight was applied on events observed on the 5’ end. Alignments with a lower score would be considered as more reliable as they would correspond to a miRNA with less editing events. The alignment score is calculated as follow:

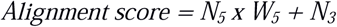

where *N_5_* and *N_3_* represent the number of editing events observed on the 5’ and 3’ extremities of the read, respectively. *W_5_* is the weight applied to the events observed on the 5’ end. Several *W_5_* values were tested and their impact on the alignment results are shown in Supplementary Table 2. Finally, for our application, *W_5_* was set to 4 in order to resolve as many ambiguous alignments as possible without penalizing too much 5’ events compared to 3’ events as they represent ~60% of aligned reads. Templated tailed nucleotides do not count as editing events as they tend to validate the parentage of a read to its reference. These templated nucleotides are most likely the result of imprecise cleavage by DROSHA and DICER. However, they might occasionally result from the action of the terminal nucleotidyl transferase that adds the same nucleotides as those surrounding the original sequence, in which case they cannot be distinguished.

By the end of the scoring algorithm, the alignment with the lowest score was retained. In case of an ambiguous read with *n* possible alignments having the same score, all alignments are kept and assigned to a weight of *1/n*, and corresponding alignments are listed in an output file (named *remaining_ambiguous.annot*).

Of note, bowtie2 also integrates an alignment score. However, this scoring is general and does not integrate the biological knowledge on editing events specific to miRNAs. The OPTIMIR scoring algorithm also differs from the one recently proposed in IsomiR-SEA pipeline (Urgese et al., 2016) which is based on the number of observed mismatches and the difference in size between a given read and the reference mature miRNAs.

#### Genotype consistency analysis

In case users provide genetic information for individuals that have been miRSeq profiled, the last step of the OPTIMIR workflow is to provide a comparison analysis of the genotype data provided by the user (in a standard vcf format) and the genotype data that could be inferred from the sequenced reads aligned to polymiRs. For an individual whose reads aligned onto a polymiR sequence that harbors the alternate allele of a SNP, consistency will be called if this individual is either heterozygous or homozygous for this allele in the provided vcf genotype file. Inconsistent alignments are discarded but saved in an output file (named *inconsistents.sam*).

Indeed, it may occur that some reads align to a polymiR sequence that harbors a given allele in an individual that is not expected to carry it. This could be due to modifications induced during library preparation or sequencing that mimic genetic variations (Wright, 2019). However, for a given individual and a given polymiR, if this event is observed for a large number of reads, another explanation must be looked for. For instance, this could occur when reads originate from sequenced fragments of other small non-coding RNAs that share a high similarity with a polymiR. Such situations are detailed in a separate output (named *consistency_table.annot*).

To detect these situations, we set up a threshold based on the number of reads aligned to each allele of the polymiR: if the presence of an alternative allele is supported by more than 1% of the total reads (i.e reads with the reference or alternative allele) that aligned to its associated polymiR, then this allele is considered as plausible. For example, given a polymiR with 980 reads aligned to the sequence integrating the reference allele and 20 reads aligned to the sequence integrating the alternative allele, then the percentage of reads supporting the presence of the alternative allele is 2%. Such a situation would then be considered inconsistent for an individual genotyped as homozygous for the reference allele and would deserve to be further investigated. Note that, when polymiRs have less than 500 aligned reads in total, a threshold of 5 supporting reads instead of a percentage of 1% was used. These parameters can be modified by the users.

Care is needed to call inconsistency when a polymiR may have homologous mature sequences and one of them is polymorphic. For example, the mature miRNA hsa-miR-1255b-5p can originate either from the pri-miRNA hsa-miR-1255b-1 located on chromosome 4 or from hsa-mir-1255b-2 located on chromosome 1. However, only the chromosome 4 copy contains a variant. If reads with the alternate allele can easily be deduced to originate from hsa-miR-1255b-1, reads with the reference allele can come from both chromosome 1 and 4 copies. As a consequence, an homozygous carriers of the alternate allele can still have reads mapping to chromosome 1 copies and such reads shall not be considered as inconsistent. We have listed in Supplementary Table 3 all mature miRNAs that have multiple pri-miRNAs sequences and tagged those that are polymorphic.

## Acknowledgments

F.T and M.R were financially supported by the GENMED Laboratory of Excellence on Medical Genomics (ANR-10-LABX-0013). DA.T was financially supported by the «EPIDEMIOM-VTE» Senior Chair from the Initiative of Excellence of the University of Bordeaux. MiRNA sequencing in the MARTHA study was performed on the iGenSeq platform (ICM Institute, Paris) and supported by a grant from the European Society of Cardiology for Medical Research Innovation.

## References

Altschul, S.F., Gish, W., Miller, W., Myers, E.W., and Lipman, D.J. (1990). Basic local alignment search tool. Journal of Molecular Biology,215, 403–410.

Ameres, S. L. and Zamore, P. D. (2013). Diversifying microRNA sequence and function. Nature Reviews. Molecular Cell Biology, 14, 475–488.

Baras, A. S., Mitchell, C. J., Myers, J. R., Gupta, S., Weng, L.-C., Ashton, J. M., Cornish, T. C., Pandey, A., and Halushka, M. K. (2015). miRge - A Multiplexed Method of Processing Small RNA-Seq Data to Determine MicroRNA Entropy. PloS One, 10:e0143066.

Bartel, D.P. (2004). MicroRNAs: genomics, biogenesis, mechanism, and function. Cell 116, 281–297.

Bofill-De Ros, X., Chen, K., Chen, S., Tesic, N., Dusan, R., Skundric, N., Nessic, S., Varjacic, V., Williams, E.H., Malhotra, R., et al. (2018). QuagmiR: A Cloud-based Application for IsomiR Big Data Analytics. Bioinformatics, https://doi.org/10.1093/bioinformatics/bty843

Burroughs, A.M., Ando, Y., de Hoon, M.J.L., Tomaru, Y., Nishibu, T., Ukekawa, R., Funakoshi, T., Kurokawa, T., Suzuki, H., Hayashizaki, Y., et al. (2010). A comprehensive survey of 3’ animal miRNA modification events and a possible role for 3’ adenylation in modulating miRNA targeting effectiveness. Genome Research, 20, 1398–1410.

Cammaerts, S., Strazisar, M., De Rijk, P., and Del Favero, J. (2015). Genetic variants in microRNA genes : impact on microRNA expression, function, and disease. Frontiers in Genetics, 6, 186.

Chen, C.J., and Heard, E. (2013). Small RNAs derived from structural non-coding RNAs. Methods, 63, 76–84.

Chiang, H.R., Schoenfeld, L.W., Ruby, J.G., Auyeung, V.C., Spies, N., Baek, D., Johnston, W.K., Russ, C., Luo, S., Babiarz, J.E., et al. (2010). Mammalian microRNAs: experimental evaluation of novel and previously annotated genes. Genes & Development, 24, 992–1009.

Danecek, P., Auton, A., Abecasis, G., Albers, C.A., Banks, E., DePristo, M.A., Handsaker, R.E., Lunter, G., Marth, G.T., Sherry, S.T., et al. (2011). The variant call format and VCFtools. Bioinformatics, 27, 2156–2158.

Desvignes, T., Loher, P., Eilbeck, K., Ma, J., Urgese, G., Fromm, B., Sydes, J., Aparicio-Puerta, E., Barrera, V., Espin, R., et al. (2018). Unification of miRNA and isomiR research: the mirGFF3 format and the mirtop API. BioRxiv doi: 10.1101/505222.

Dick, K.J., Nelson, C.P., Tsaprouni, L., Sandling, J.K., Aïssi, D., Wahl, S., Meduri, E., Morange, P.-E., Gagnon, F., Grallert, H., et al. (2014). DNA methylation and body-mass index: a genome-wide analysis. Lancet London England, 383, 1990–1998.

Duan, R., Pak, C., and Jin, P. (2007). Single nucleotide polymorphism associated with mature miR-125a alters the processing of pri-miRNA. Human Molecular Genetics, 16, 1124–1131.

Filipowicz, W., Bhattacharyya, S. N., and Sonenberg, N. (2008). Mechanisms of post-transcriptional regulation by microRNAs : are the answers in sight? Nature Reviews. Genetics, 9, 102–114.

Germain, M., Chasman, D.I., de Haan, H., Tang, W., Lindström, S., Weng, L.-C., de Andrade, M., de Visser, M.C.H., Wiggins, K.L., Suchon, P., et al. (2015). Meta-analysis of 65,734 individuals identifies TSPAN15 and SLC44A2 as two susceptibility loci for venous thromboembolism. American Journalof Human Genetics, 96, 532–542.

Git, A., Dvinge, H., Salmon-Divon, M., Osborne, M., Kutter, C., Hadfield, J., Bertone, P., and Caldas, C. (2010). Systematic comparison of microarray profiling, real-time PCR, and next-generation sequencing technologies for measuring differential microRNA expression. RNA, 16, 991–1006.

Gong, J., Tong, Y., Zhang, H.-M., Wang, K., Hu, T., Shan, G., Sun, J., and Guo, A.-Y. (2012). Genome-wide identification of SNPs in microRNA genes and the SNP effects on microRNA target binding and biogenesis. Human Mutation, 33, 254–263.

Han, M. and Zheng, Y. (2013). Comprehensive analysis of single nucleotide polymorphisms in human microRNAs. PloS One, 8 :e78028.

Heintz-Buschart, A., Yusuf, D., Kaysen, A., Etheridge, A., Fritz, J.V., May, P., de Beaufort, C., Upadhyaya, B.B., Ghosal, A., Galas, D.J., et al. (2018). Small RNA profiling of low biomass samples: identification and removal of contaminants. BMC Biology, 16, 52.

Katoh T, Hojo H, Suzuki T. (2015). Destabilization of microRNAs in human cells by 3′ deadenylation mediated by PARN and CUGBP1. Nucleic Acids Research, 43, 7521–7534.

Kawahara, Y., Zinshteyn, B., Sethupathy, P., Iizasa, H., Hatzigeorgiou, A. G., & Nishikura, K. (2007). Redirection of Silencing Targets by Adenosine-to-Inosine Editing of miRNAs. Science (New York, N.Y.), 315, 1137–1140.

Kawamata, T., & Tomari, Y. (2010). Making RISC. Trends in Biochemical Sciences, 35, 368–376.

Kim, V. N., Han, J., and Siomi, M. C. (2009). Biogenesis of small RNAs in animals. Nature Reviews Molecular Cell Biology, 10, 126–139.

Kim, Y.-K., Kim, B., and Kim, V. N. (2016). Re-evaluation of the roles of DROSHA, Exportin 5, and DICER in microRNA biogenesis. Proceedings of the National Academy of Sciences of the United States of America, 113, E1881–E1889.

Kozomara, A. and Griffiths-Jones, S. (2014). miRBase : annotating high confidence microRNAs using deep sequencing data. Nucleic Acids Research, 42, D68–D73.

Langmead, B. and Salzberg, S. L. (2012). Fast gapped-read alignment with Bowtie 2. Nature Methods, 9, 357.

Lorenz, R., Bernhart, S. H., Höner zu Siederdissen, C., Tafer, H., Flamm, C., Stadler, P. F., and Hofacker, I. L. (2011). ViennaRNA Package 2.0. Algorithms for Molecular Biology, 6, 26.

Martin, M. (2011). Cutadapt removes adapter sequences from high-throughput sequencing reads. EMBnet.journal, 17, 10–12.

Meijer, H.A., Smith, E.M., and Bushell, M. (2014). Regulation of miRNA strand selection: follow the leader? Biochemical Society Transactions, 42, 1135–1140.

Mencía, A., Modamio-Høybjør, S., Redshaw, N., Morín, M., Mayo-Merino, F., Olavarrieta, L., Aguirre, L. A., del Castillo, I., Steel, K. P., Dalmay, T., Moreno, F., and Moreno-Pelayo, M. A. (2009). Mutations in the seed region of human miR-96 are responsible for nonsyndromic progressive hearing loss. Nature Genetics, 41, 609–613.

Mitchell, P.S., Parkin, R.K., Kroh, E.M., Fritz, B.R., Wyman, S.K., Pogosova-Agadjanyan, E.L., Peterson, A., Noteboom, J., O’Briant, K.C., Allen, A., et al. (2008). Circulating microRNAs as stable blood-based markers for cancer detection. Proceedings of the National Academy of Science, 105, 10513–10518.

Morales, S., Monzo, M., and Navarro, A. (2017). Epigenetic regulation mechanisms of microRNA expression. Biomolecular Concepts, 8, 203–212.

Neilsen, C. T., Goodall, G. J., and Bracken, C. P. (2012). IsomiRs–the overlooked repertoire in the dynamic microRNAome. Trends in genetics, 28, 544–549.

Nishikura, K. (2016). A-to-I editing of coding and non-coding RNAs by ADARs. Nature Reviews Molecular Cell Biology, 17, 83–96.

Oudot-Mellakh, T., Cohen, W., Germain, M., Saut, N., Kallel, C., Zelenika, D., Lathrop, M., Trégouët, D.- A., and Morange, P.-E. (2012). Genome wide association study for plasma levels of natural anticoagulant inhibitors and protein C anticoagulant pathway : the MARTHA project. British Journal of Haematology, 157, 230–239.

Pulcrano-Nicolas, A.-S., Proust, C., Clarençon, F., Jacquens, A., Perret, C., Roux, M., Shotar, E., Thibord, F., Puybasset, L., Garnier, S., et al. (2018). Whole-Blood miRNA Sequencing Profiling for Vasospasm in Patients With Aneurysmal Subarachnoid Hemorrhage. Stroke, 49, 2220–2223.

Roux, M., Perret, C., Feigerlova, E., Mohand Oumoussa, B., Saulnier, P.-J., Proust, C., Trégouët, D.-A., and Hadjadj, S. (2018). Plasma levels of hsa-miR-152-3p are associated with diabetic nephropathy in patients with type 2 diabetes. Nephrology, Dialysis, Transplantation: Official Publication of the European Dialysis and Transplant Association - European Renal Association. https://doi.org/10.1093/ndt/gfx367.

Russell, P.H., Vestal, B., Shi, W., Rudra, P.D., Dowell, R., Radcliffe, R., Saba, L., and Kechris, K. (2018). miR-MaGiC improves quantification accuracy for small RNA-seq. BMC Research Notes, 11, 296.

Tam, S., de Borja, R., Tsao, M.-S., and McPherson, J. D. (2014). Robust global microRNA expression profiling using next-generation sequencing technologies. Laboratory Investigation; a Journal of Technical Methods and Pathology, 94, 350–358.

Urgese, G., Paciello, G., Acquaviva, A., & Ficarra, E. (2016). isomiR-SEA: an RNA-Seq analysis tool for miRNAs/isomiRs expression level profiling and miRNA-mRNA interaction sites evaluation. BMC Bioinformatics, 17, 148.

Wallaert, A., Van Loocke, W., Hernandez, L., Taghon, T., Speleman, F., and Van Vlierberghe, P. (2017). Comprehensive miRNA expression profiling in human T-cell acute lymphoblastic leukemia by small RNA-sequencing. Scientific Reports, 7, 7901.

Wright, C., Rajpurohit, A., Burke, E.E., Williams, C., Collado-Torres, L., Kimos, M., Brandon, N.J., Cross, A.J., Jaffe, A.E., Weinberger, D.R., et al. (2019). Comprehensive assessment of multiple biases in small RNA sequencing reveals significant differences in the performance of widely used methods. BioRxiv doi: 10.1101/445437.

Wu, X., Kim, T.-K., Baxter, D., Scherler, K., Gordon, A., Fong, O., Etheridge, A., Galas, D. J., and Wang, K. (2017). sRNAnalyzer-a flexible and customizable small RNA sequencing data analysis pipeline. Nucleic Acids Research, 45, 12140–12151.

Wu, C.W., Evans, J.M., Huang, S., Mahoney, D.W., Dukek, B.A., Taylor, W.R., Yab, T.C., Smyrk, T.C., Jen, J., Kisiel, J.B., et al. (2018). A Comprehensive Approach to Sequence-oriented IsomiR annotation (CASMIR): demonstration with IsomiR profiling in colorectal neoplasia. BMC Genomics, 19, 401.

Wyman, S. K., Knouf, E. C., Parkin, R. K., Fritz, B. R., Lin, D. W., Dennis, L. M., Krouse, M. A., Webster, P. J., and Tewari, M. (2011). Post-transcriptional generation of miRNA variants by multiple nucleotidyl transferases contributes to miRNA transcriptome complexity. Genome Research, 21, 1450–1461.

Ziemann, M., Kaspi, A., and El-Osta, A. (2016). Evaluation of microRNA alignment techniques. RNA, 22, 1120–1138.

