## Supplemental Figures 1 & 2 for "OPTIMIR, a novel algorithm for integrating available genome-wide genotype data into miRNA sequence alignment analysis"

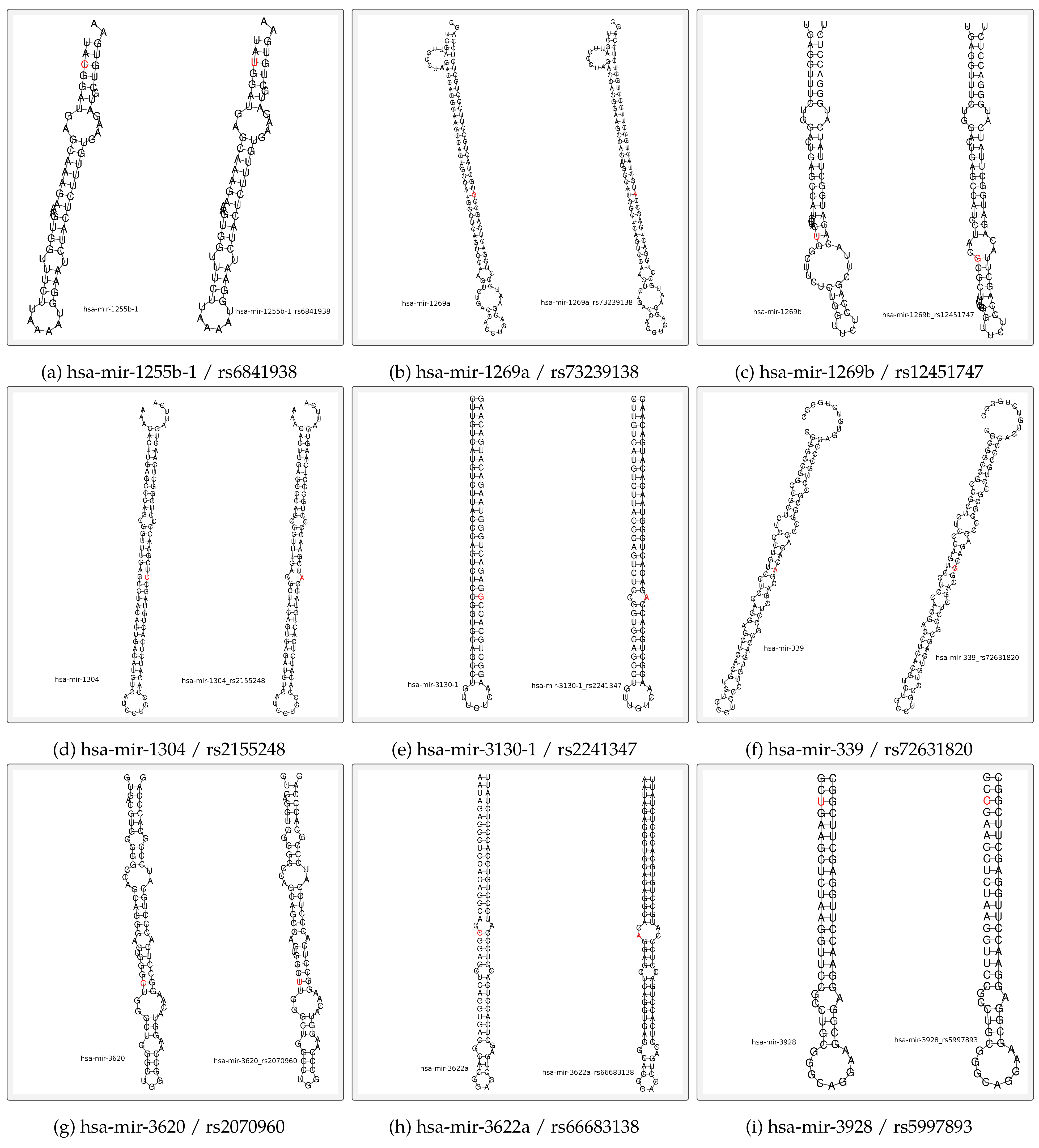
**Supplementary Figure S1 (1/4): RNAfold predictions of polymiRs secondary structures that are expressed by heterozygous samples**

**Supplementary Figure S1 (2/4): RNAfold predictions of polymiRs secondary structures that are expressed by heterozygous samples**

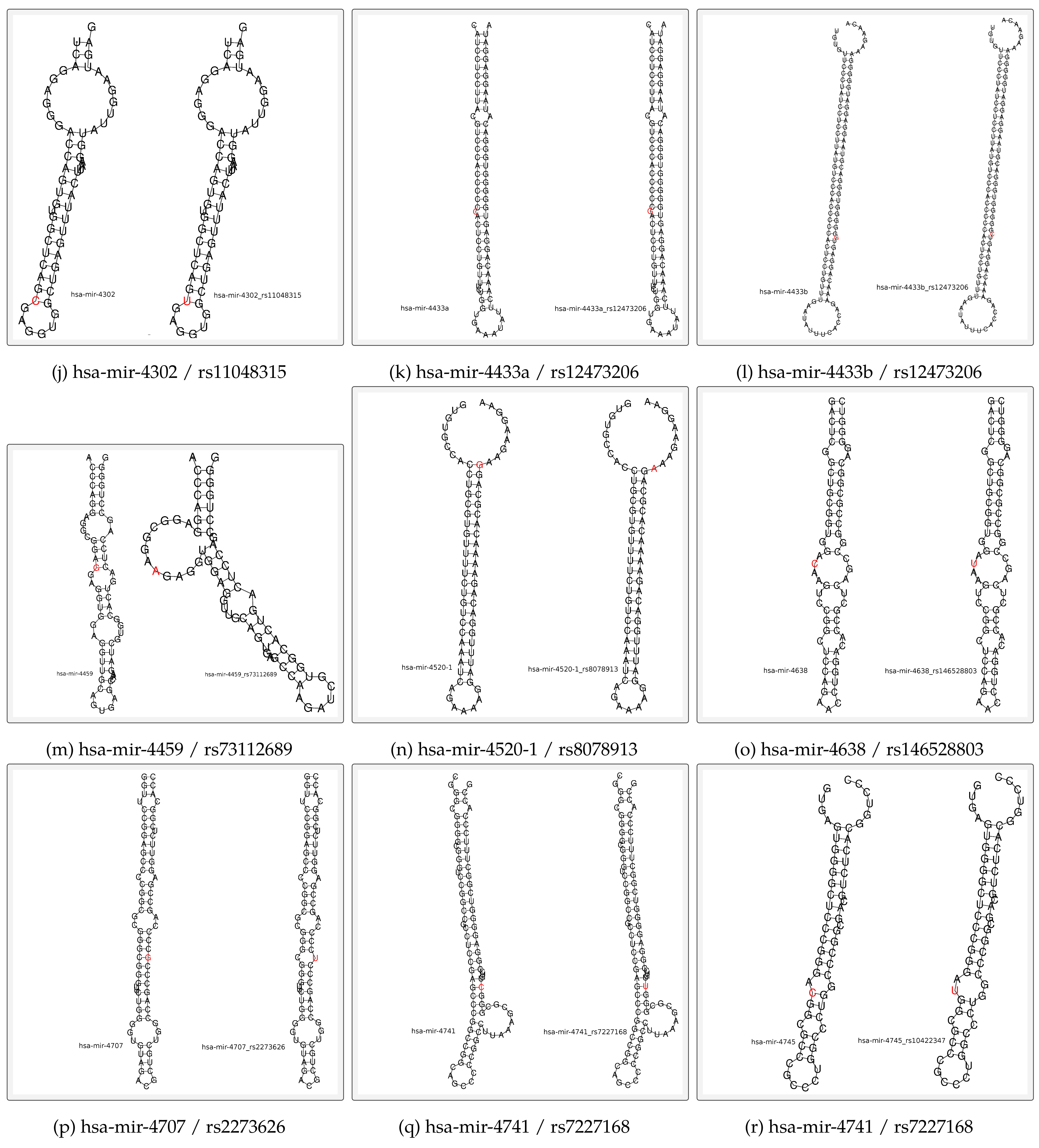

**Supplementary Figure S1 (3/4): RNAfold predictions of polymiRs secondary structures that are expressed by heterozygous samples**

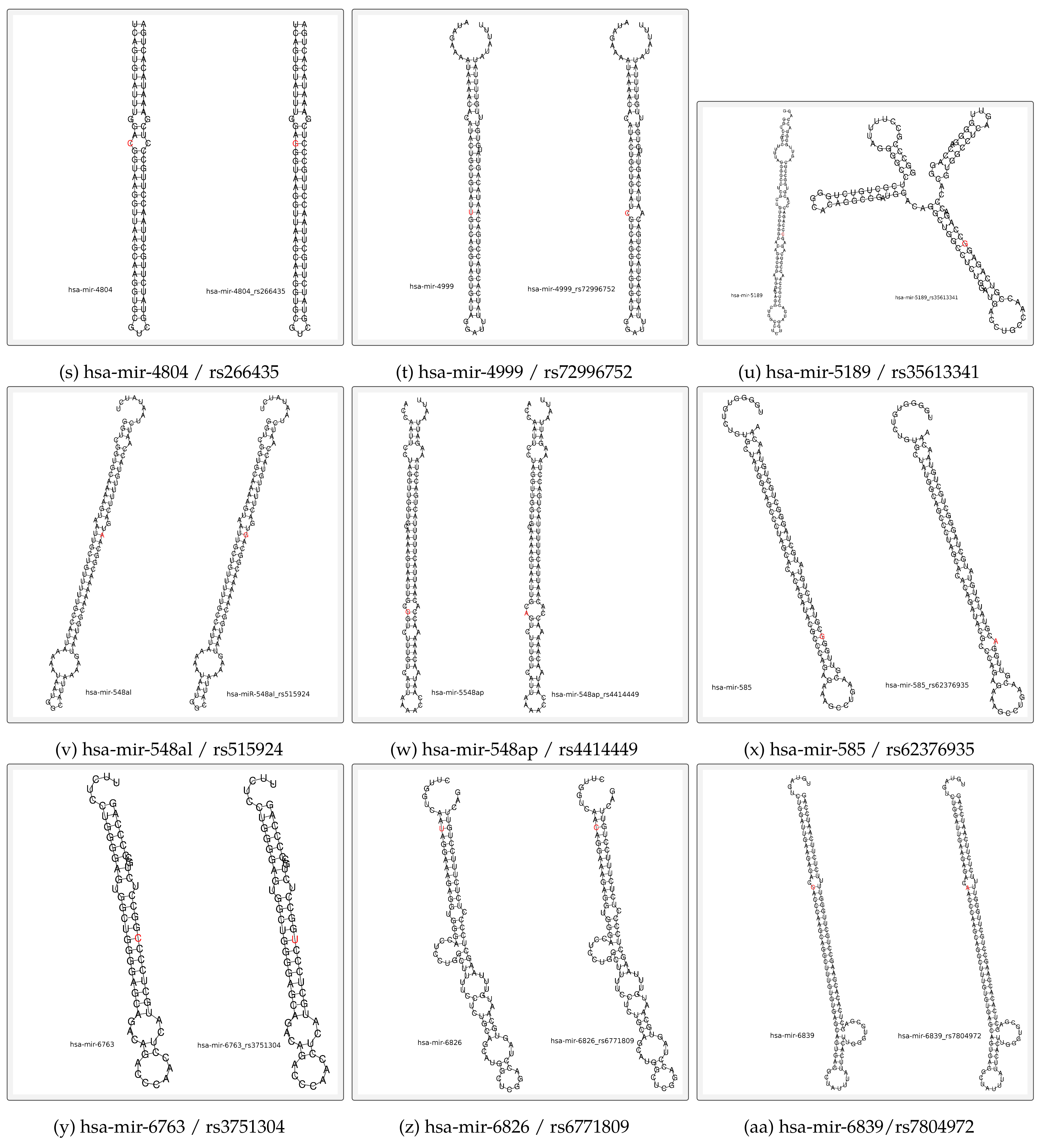

**Su****pplementary Figure S1 (4/4): RNAfold predictions of polymiRs secondary structures that are expressed by heterozygous samples**

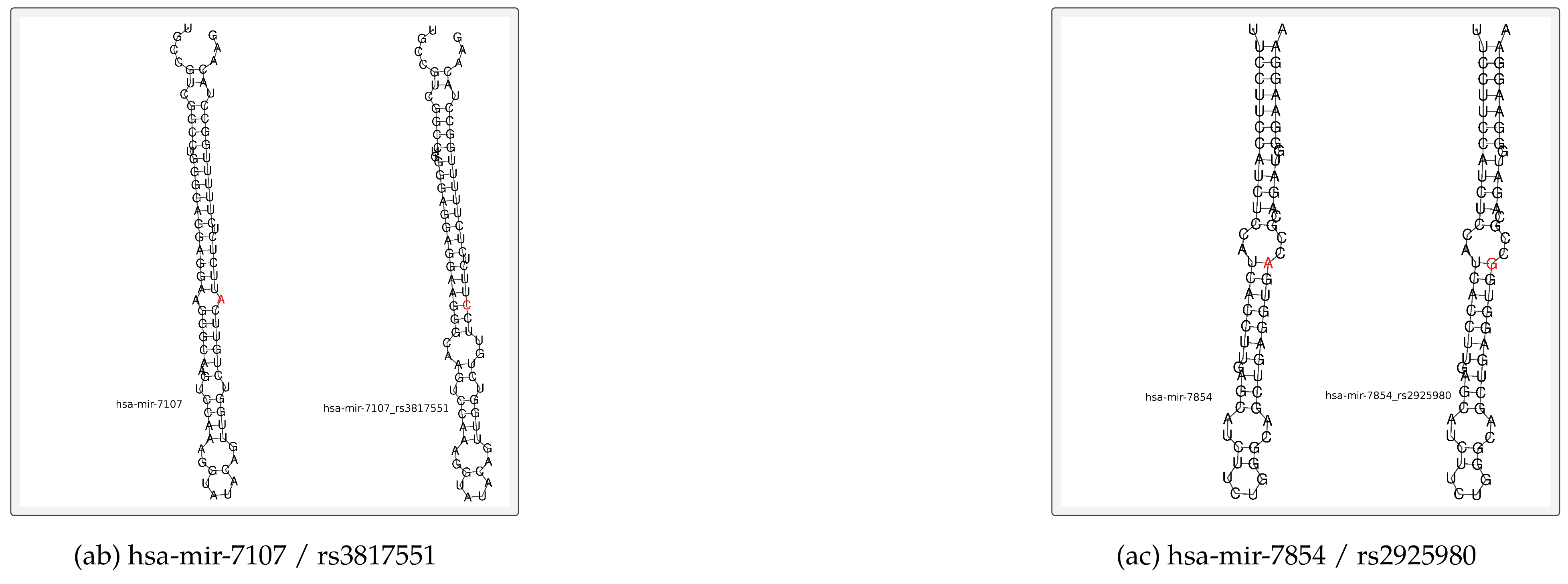

**Supplementary Figure S2 : Library preparation workflow**

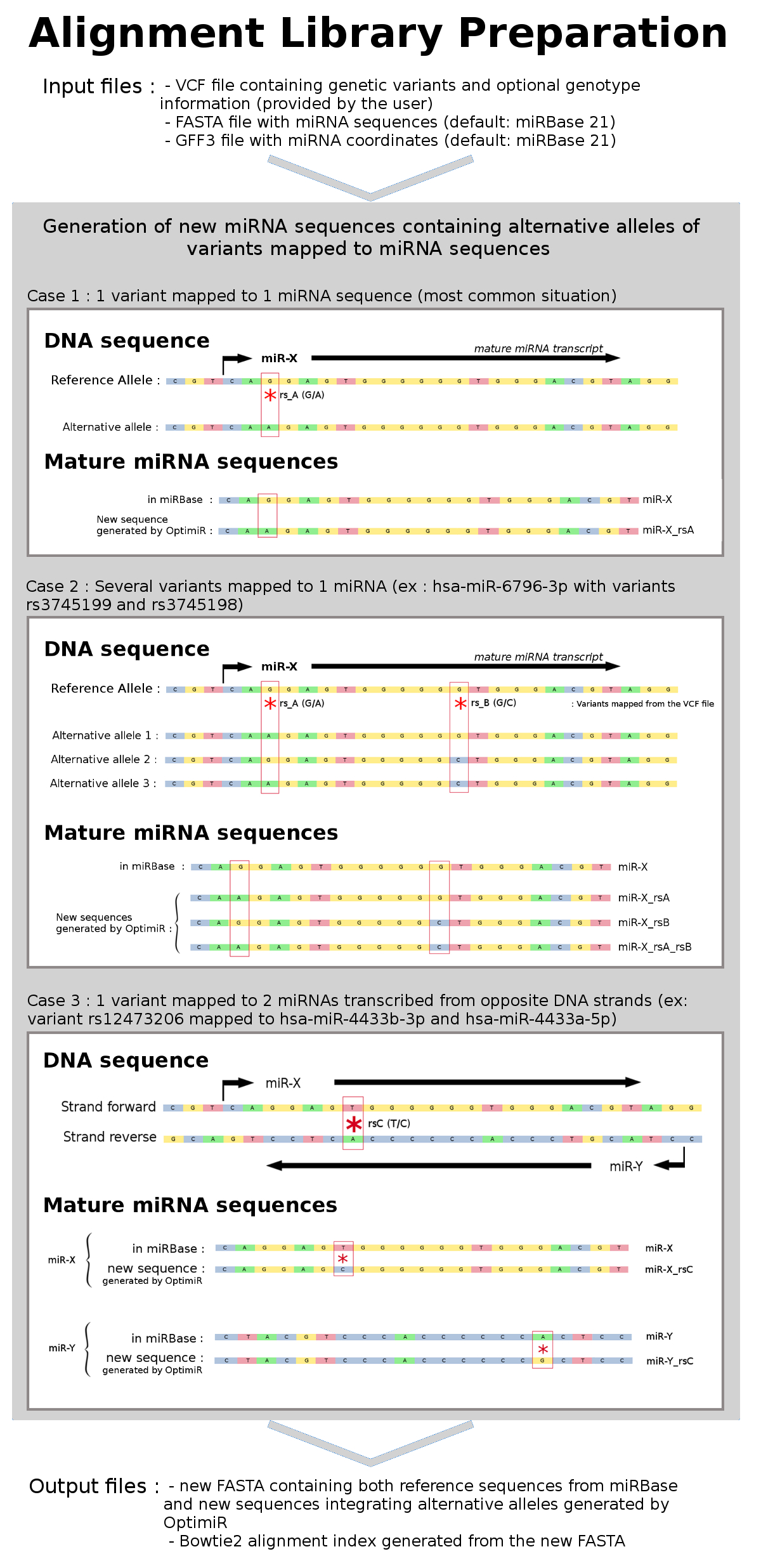
